# Comparison of Anatomical and Diffusion MRI for detecting Parkinson’s Disease using Deep Convolutional Neural Network

**DOI:** 10.1101/2023.05.01.538952

**Authors:** Tamoghna Chattopadhyay, Amit Singh, Emily Laltoo, Christina P. Boyle, Conor Owens-Walton, Yao-Liang Chen, Philip Cook, Corey McMillan, Chih-Chien Tsai, J-J Wang, Yih-Ru Wu, Ysbrand van der Werf, Paul M. Thompson

## Abstract

Parkinson’s disease (PD) is a progressive neurodegenerative disease that affects over 10 million people worldwide. Brain atrophy and microstructural abnormalities tend to be more subtle in PD than in other age-related conditions such as Alzheimer’s disease, so there is interest in how well machine learning methods can detect PD in radiological scans. Deep learning models based on convolutional neural networks (CNNs) can automatically distil diagnostically useful features from raw MRI scans, but most CNN-based deep learning models have only been tested on T1-weighted brain MRI. Here we examine the added value of diffusion-weighted MRI (dMRI) - a variant of MRI, sensitive to microstructural tissue properties - as an additional input in CNN-based models for PD classification. Our evaluations used data from 3 separate cohorts - from Chang Gung University, the University of Pennsylvania, and the PPMI dataset. We trained CNNs on various combinations of these cohorts to find the best predictive model. Although tests on more diverse data are warranted, deep-learned models from dMRI show promise for PD classification.

**Clinical Relevance:** This study supports the use of diffusion-weighted images as an alternative to anatomical images for AI-based detection of Parkinson’s disease.

## I. INTRODUCTION

Parkinson’s disease (PD) is the second most common neurological disorder worldwide, affecting over 10 million people, and its prevalence is increasing faster than any other neurological disorder [1]. PD is characterized by cardinal motor symptoms including tremor at rest, rigidity and postural instability [2]. Neuropathological abnormalities are found at autopsy in PD, including α-synuclein-immunopositive Lewy bodies and neurites [3], which are associated with loss of dopaminergic neurons in the *substantia nigra pars compacta*, leading to dopamine depletion in the striatum [4]. Large-scale neuroimaging studies have begun to identify robust stage-specific structural brain abnormalities in PD, yielding *in vivo* signatures of the disease process that mirror the anatomic progression of neuropathology [5]. Improved biomarkers may assist diagnosis, staging and prognosis in PD, which now rely primarily on clinical evaluations of motor and non-motor impairments. Machine learning has been applied to structural brain MRI to discover metrics that best differentiate PD participants from healthy controls, with wide variation in performance (for a review, see [6]). As white matter microstructure is also impaired in PD [7], diffusion tensor imaging (DTI) may improve these models, as abnormal DTI metrics have been reported in PD prior to the appearance of structural brain atrophy [8].

Here we set out to classify individuals with PD versus healthy controls, based on 3D convolutional neural networks trained on both anatomical and diffusion MRI. CNNs are attractive, as they can learn predictive features from raw images. Diffusion-weighted brain MRI (dMRI) is sensitive to subtle alterations in the brain’s microstructure and can offer independent information on white matter integrity that is not detectable with standard T1-weighted MRI. Here we tested the performance of 3D CNNs for classifying PD based on T1w MRI, as well as DTI-derived maps of mean, radial and axial diffusivity (MD/RD/AD) and fractional anisotropy (FA). We also tested the performance when combining the two multimodal data types in concatenated CNN architectures.

## II. DATA

Data for this project was drawn from 3 cohorts (**Table 1**). The T1-weighted brain MRI volumes were pre-processed using a sequence of steps, including nonparametric intensity normalization (N4 bias field correction) [9], ‘skull-stripping’ for brain extraction, registration to a template with 6 degrees of freedom (rigid-body) registration and isometric voxel resampling to 2 mm. The pre-processed images were of size 91×109×91. The T1w images were scaled to take values between 0 and 1 via min-max scaling. DTI images were pre-processed using the ENIGMA-DTI Pipeline, which includes image denoising, Gibbs de-ringing, eddy current correction, echo-planar imaging (EPI)-induced distortion correction and bias-field correction. This process produced a DTI map of the FA, MD, RD and AD, with each map registered to the pre-processed T1 space.

**TABLE 1.** Demographics of the Datasets used for Training and Testing. PD/CTL denotes the numbers of PD patients and controls. H&Y stage denotes the Hoehn & Yahr stage - a commonly used PD staging system where 1 denotes the most mildly impaired, and 5 the most severely impaired.

| Dataset | N | Age<br>(mean±<br>SD) | Sex<br>(F/M) | PD/<br>CTL | Mean<br>H&Y<br>Stage |
| --- | --- | --- | --- | --- | --- |
| Taiwan | 529 | 60.6 ± 8.9 | 254 /<br>275 | 306 /<br>223 | 2.24 |
| UPenn | 165 | 66.7 ± 7.9 | 63 /<br>105 | 138 /<br>30 | 2.64 |
| PPMI | 183 | 62.7 ± 9.7 | 66 /<br>117 | 129 /<br>54 | 1.59 |

**TABLE II.** Results for all experiments, showing balanced accuracy and F1 Scores for all modalities. The highest values are in **bold**.

|  |  | T1w |  | DWI-FA |  | DWI-MD |  | DWI-RD |  | DWI-AD |  |
| --- | --- | --- | --- | --- | --- | --- | --- | --- | --- | --- | --- |
|  |  | Balanced Accuracy | F1 score | Balanced Accuracy | F1 score | Balanced Accuracy | F1 score | Balanced Accuracy | F1 score | Balanced Accuracy | F1 score |
| Experiment 1 | Test set | 0.537 | 0.797 | 0.708 | 0.826 | 0.796 | 0.851 | 0.793 | 0.847 | **0.803** | **0.860** |
|  | PPMI | 0.494 | 0.805 | 0.535 | 0.822 | 0.555 | 0.778 | 0.524 | 0.771 | **0.567** | **0.754** |
| Experiment 2 | Test set | 0.500 | 0.759 | 0.548 | 0.744 | 0.683 | 0.797 | **0.726** | **0.810** | 0.639 | 0.787 |
|  | UPenn | 0.500 | 0.902 | 0.486 | 0.852 | 0.578 | 0.782 | 0.540 | 0.872 | **0.577** | **0.882** |
| Experiment 3 | Test set | 0.566 | 0.706 | 0.717 | 0.781 | 0.736 | 0.796 | **0.747** | 0.797 | 0.728 | **0.799** |
| Experiment 4 | Test set | 0.500 | 0.792 | 0.608 | 0.816 | **0.704** | **0.814** | 0.666 | 0.802 | 0.651 | 0.799 |

**TABLE III.**
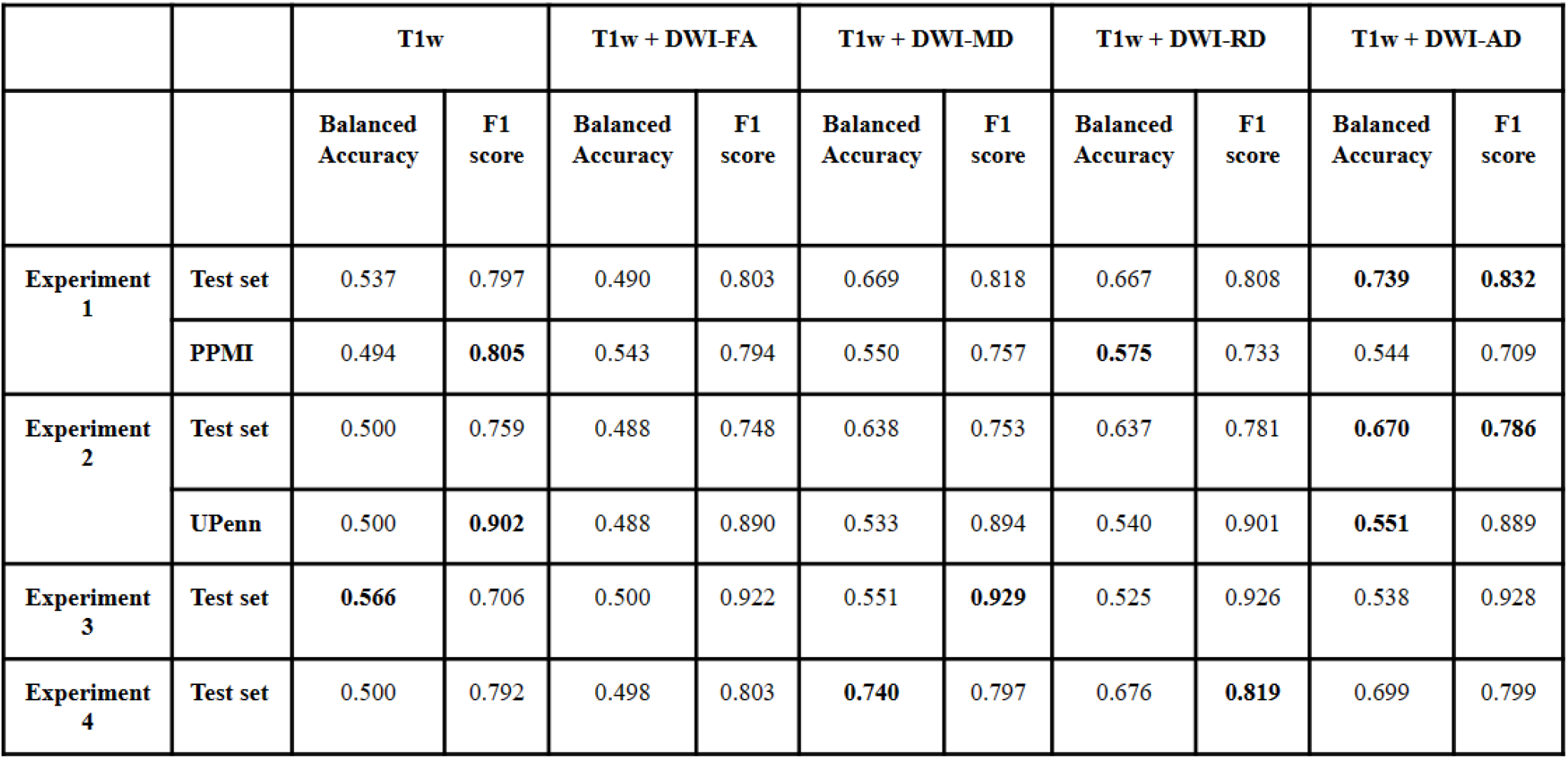
Results for all experiments, showing balanced accuracy and F1 Scores for all pairs of combined modalities. The highest values are in **bold**.

## III. MODEL AND METHODS

After registering the DWI maps and T1s to a common template, we used elastic deformation, a technique widely used in medical image processing, to augment the training data. We used displacement vectors and a spline interpolation for input image deformation. The 3D CNN architecture (**Figure 1**) consisted of four 3D convolution layers with a 3×3 filter size, followed by one 3D convolution layer with a 1×1 filter, and a final Dense layer with a sigmoid activation function. All layers used the ReLu activation function and Instance Normalization. Dropout layers, with a dropout rate of 0.5, and a 3D Average Pooling layer with a 2×2 filter size were added to the 2^nd^, 3^rd^, and 4^th^ layers. Models were trained with a learning rate of 1e-4, and test performance was assessed using balanced accuracy. To deal with overfitting, both L1 and L2 regularizers were used, along with dropouts between layers and early stopping. Hyperparameter tuning was performed by running *k*-fold cross validation.

**Fig. 1.**
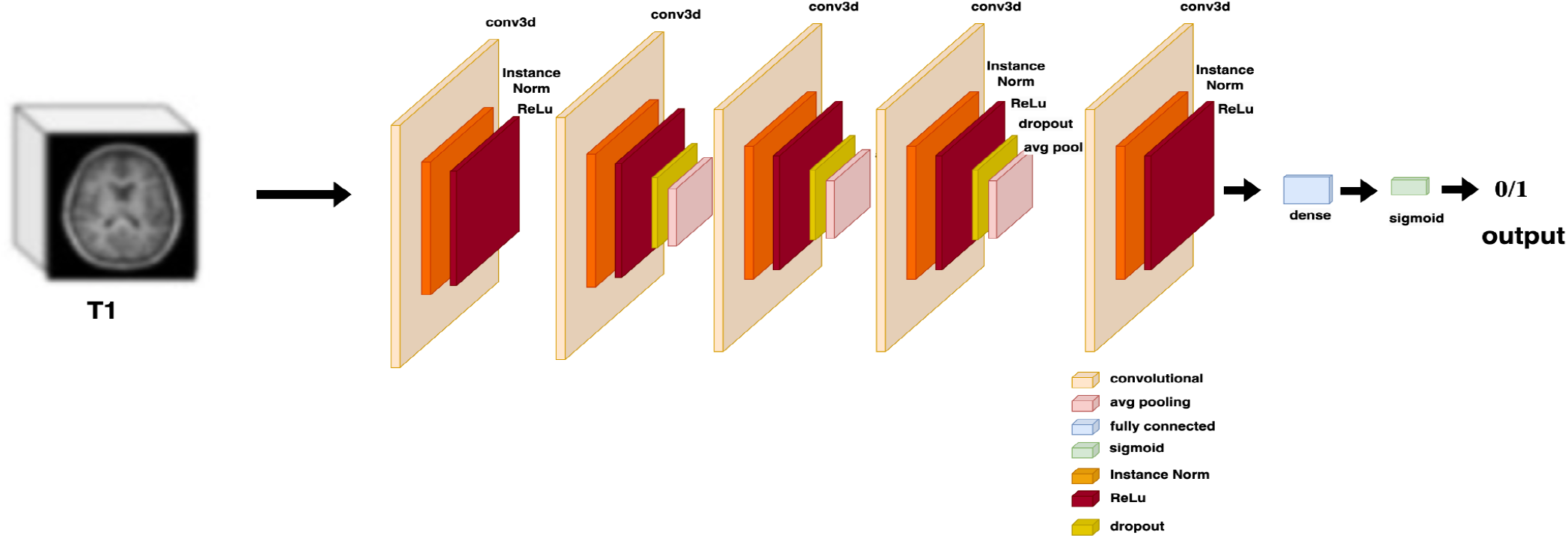
3D CNN Architecture used for training on data from a single modality (here, anatomical MRI).

Four sets of experiments were run using the 3D CNN architecture. In the first experiment, the training, validation and test dataset for the model was a combination of the Taiwan and UPenn datasets, while the PPMI dataset was kept as a holdout test set. The Taiwan and UPenn datasets were divided in the ratio 80:10:10 for training (n=557), validation (n=70) and testing (n=70). In the second experiment, the training, validation and test dataset for the model was a combination of the Taiwan and PPMI datasets, while the UPenn dataset was held out as a test set. The Taiwan and PPMI datasets were divided in the ratio 80:10:10 for training (n=568), validation (n=72) and testing (n=72).

In the third experiment, all datasets were combined and divided into training (n=617), validation (n=175) and testing (n=90) in the ratio 70:20:10. In the fourth experiment, the datasets were combined in the ratio 80:10:10 individually to create training (n=702), validation (n=89) and test (n=89) sets. The same model was used for training in all four experiments. The 3D-CNN was trained for 100 epochs with the Adam (with weight decay) optimizer. In all four experiments, the batch size was set to 8 and the model was trained until the validation loss did not improve for 10 consecutive epochs. Each experiment was performed three times, and the average values of balanced accuracy and F1 scores were retained to compare the performance.

For the concatenation, we reworked the above architecture. The 3D CNN architecture (**Figure 2**) consisted of the same four 3D Convolution layers, but after flattening, they were concatenated and sent through a Dense Layer with sigmoid activation function. This Y-shaped architecture uses separate CNNs to distil predictive features from the anatomical MRI and diffusion MRI, which are then merged for disease classification. The same four experiments, as in the case of single modality, were repeated for the combination of images. The 3D-CNN was trained for 100 epochs with Adam (with weight decay) optimizer, batch size was set to 4. The model was trained until validation loss did not improve for 10 consecutive epochs. Each experiment was performed three times, and average values of balanced accuracy and F1 scores were retained to compare the performance.

**Fig. 2.**
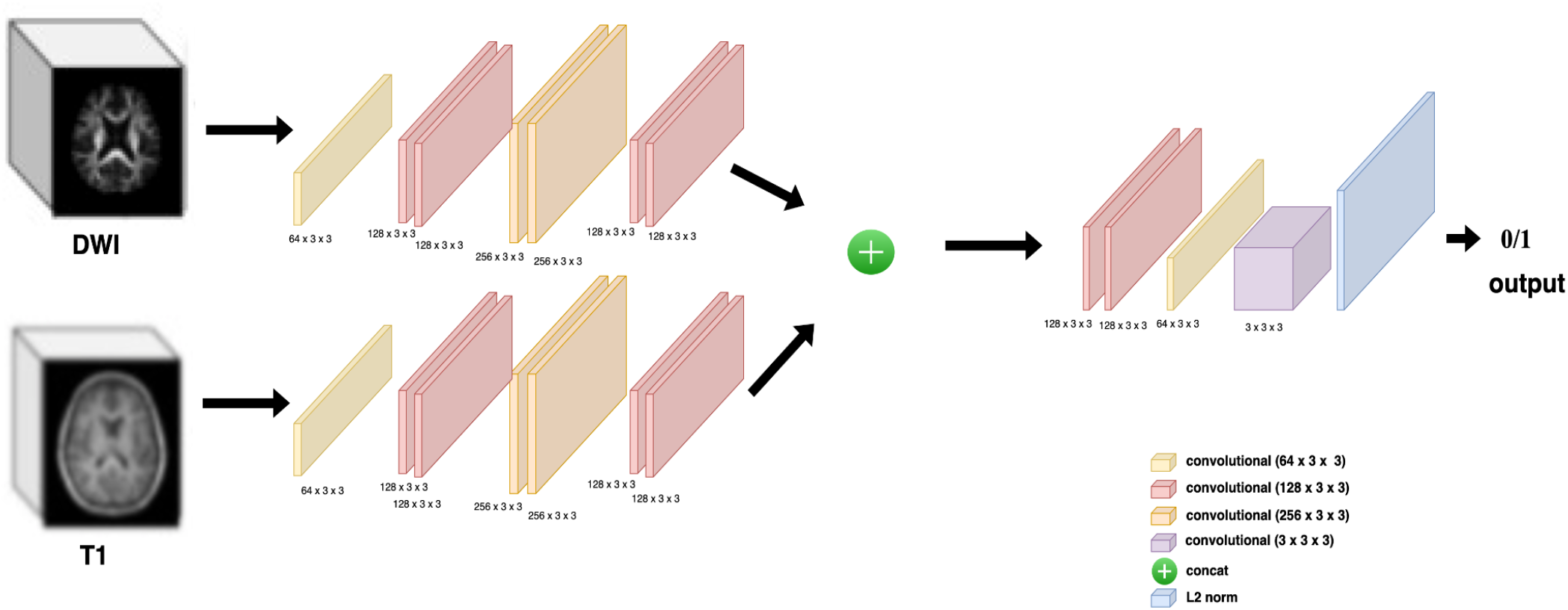
3D CNN Architecture used for dual modality training.

So, for the first experiment the training data was a combination of Taiwan and UPenn dataset, with PPMI as holdout Test dataset. The second experiment had a combination of Taiwan and PPMI as training and UPenn as a holdout test set. The other two experiments used combinations of all three datasets in different proportions.

## IV. RESULTS

Among the single modality experiments, for the first experiment, the best balanced accuracy was 0.803 for DWI-AD with an F1 score of 0.860. This model gave a balanced accuracy of 0.567 and an F1 score of 0.754 on the hold-out PPMI dataset. The worst balanced accuracy was 0.537 for T1w with an F1 score of 0.797. This model gave a balanced accuracy of 0.494 and an F1 score of 0.805 on the hold-out PPMI dataset. For the second experiment, the best balanced accuracy was 0.726 for DWI-RD with an F1 score of 0.810. This model gave a balanced accuracy of 0.540 and an F1 score of 0.872 on the hold-out UPenn dataset. The worst balanced accuracy was 0.500 for T1w with an F1 score of 0.759. This model gave a balanced accuracy of 0.500 and an F1 score of 0.902 on the held-out UPenn dataset. For the third experiment, the best balanced accuracy was 0.747 for DWI-RD with a F1 score of 0.797. The worst balanced accuracy was 0.566 for T1w with an F1 score of 0.706. For the fourth experiment, the best balanced accuracy was 0.704 for DWI-MD with an F1 score of 0.814. The worst balanced accuracy was 0.500 for T1w with an F1 score of 0.792. Based on the balanced accuracy, the diffusion MRI maps always outperformed the T1w as input data. The RoC-AUC Curves for all experiments are shown in **Figure 3-6**. As these figures show, the area under the curve is worst for T1ws as input in all experiments.

**Fig 3.**
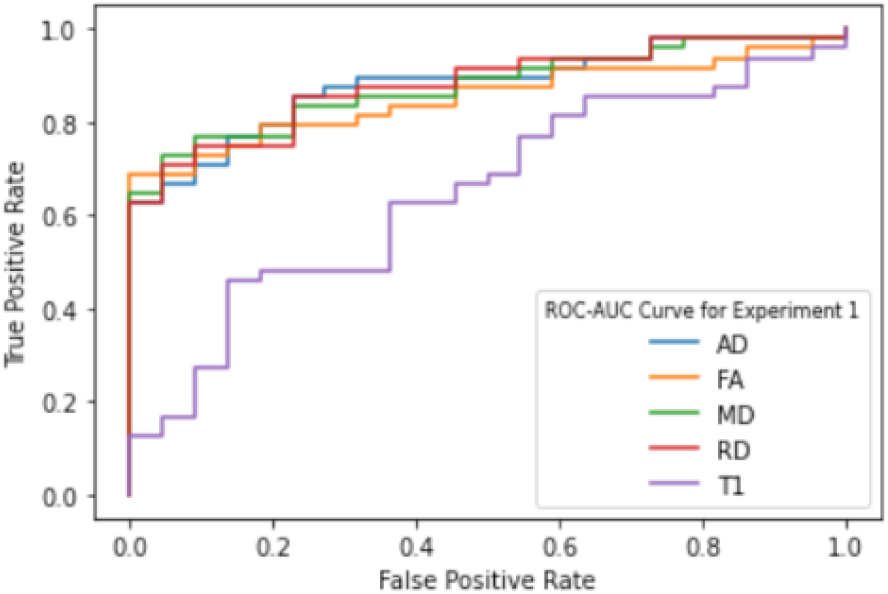
ROC-AUC Curve for Experiment 1

**Fig 4.**
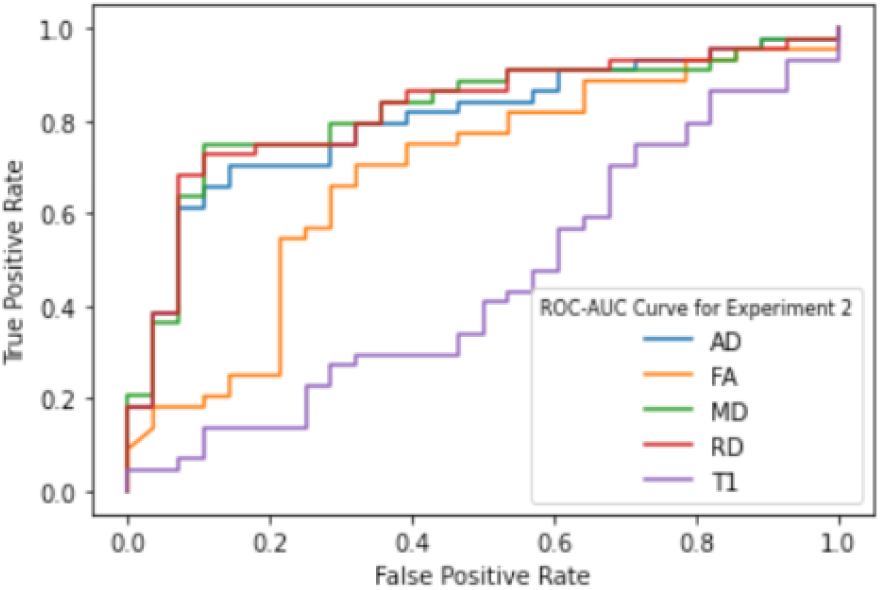
ROC-AUC Curve for Experiment 2

**Fig 5.**
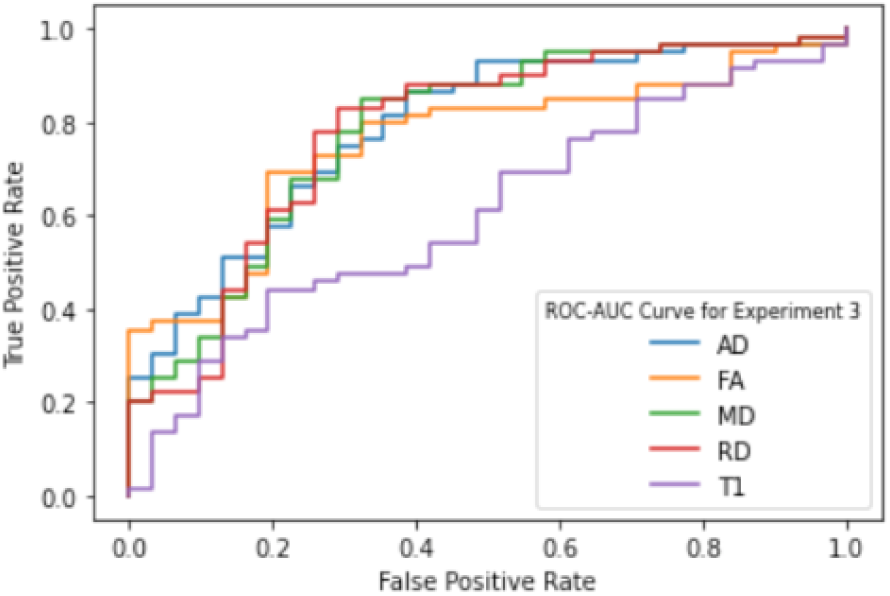
ROC-AUC Curve for Experiment 3

**Fig 6.**
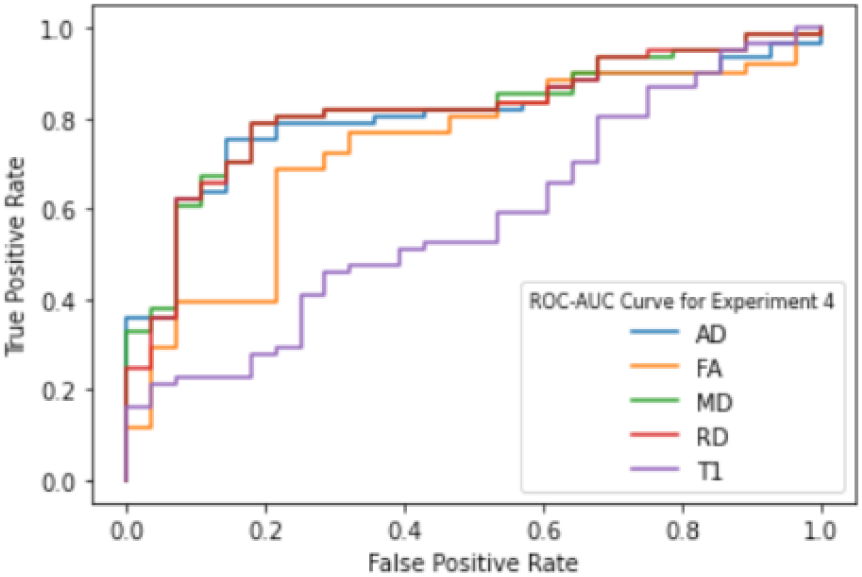
ROC-AUC Curve for Experiment 4

An independent samples *t*-test found that the AUC of participants in the T1w group was significantly lower than the AUC when using DWI images in all four experiments. For experiment 1, t(10) = -14.9 and *p*<0.0001. For experiment 2, *t*(8) = -84.6 and *p*<0.0001. For experiment 3, *t*(7.1) = -6.0 and *p* = 0.0002. For experiment 4, t(9.7) = -14.2 and *p*<0.0001. Thus, in general from our experiments, DTI measures outperformed T1w for PD classification, and there was some evidence that the diffusivity measures outperformed FA, but additional independent data is needed to verify this.

For the dual modality experiments, combining T1w and DWI-MD and DWI-AD gave the best results, compared to the other two combinations. In most cases, the balanced accuracy increases when a combination of T1w and dMRI was used, relative to using T1 alone. Even so, more data may be required to improve the accuracy of these models as the dimension of input data is large and the amount of training data is small: with the current amount of training data, the balanced accuracy of the fused model is still less than the case where DWIs are the only input; intuitively, the fused model should be better, as it uses more information, so long as there is sufficient data to train it. Another interesting conclusion was that PPMI was much harder to classify as compared to the other two datasets. This may be because the mean H&Y stage of the patients in the three datasets is in the following order: UPenn (2.64) > Taiwan (2.24) > PPMI (1.59), so PPMI patients may generally have milder brain abnormalities that those in the other datasets.

## V. CONCLUSION AND DISCUSSION

In this work, we trained deep learning models to classify individuals as PD patients or healthy controls, using 3D CNNs and different types of brain MRI. In a novel approach, building on [13], we tested diffusion MRI maps as inputs and, with the methods tested, they outperformed standard anatomical MRI. While prior studies have found both structural and diffusion MRI abnormalities in PD, the explicit segmentation of regions of interest often requires time-consuming quality control and human interaction with the data, making it hard to design a practical classifier from these features. We explored various combinations of the three cohorts of data, to find the best model to generalize over new data. The best accuracy was around 0.75 for DWI-AD, which offers a promising baseline for further testing.

For combined modalities, due to the increase in the number of parameters, more data is required for training and improving the performance. Another interesting conclusion was that given the mean H&Y stage of the three datasets is UPenn (2.64) > Taiwan (2.24) > PPMI (1.59), PPMI is much harder to classify, compared to the other two datasets.

## VI. FUTURE WORK

Future work will train and test the methods on more diverse datasets, and will assess how well the models generalize to patients at different stages of PD, and with mixed diagnoses. We will test the added value of training classifiers on more MRI and DTI data, along with data modalities that are less commonly collected, such as quantitative parametric MRI, DAT-SPECT, and resting state fMRI, all of which may offer complementary information for PD classification. We will examine other training techniques such as transfer learning, and other CNN variants such as DenseNet-121, which has more depth than a standard 3D CNN. A strong PD classifier may also serve as a guide to creating deep learning methods for more challenging tasks, such as PD staging [10], differential diagnosis and subtyping, predicting future decline, and predicting response to treatment or other interventions.

## VII. COMPLIANCE WITH ETHICAL STANDARDS

We obtained IRB approval for this analysis of previously collected, anonymized de-identified data. We also analyzed anonymized human subject data made available in open access by PPMI.

## ACKNOWLEDGMENTS

This work was supported by NINDS grant R01NS107513.

